# Enhanced diastolic dysfunction but preserved systolic function after acute pressure overload in the absence of smoothelin-like 1 protein

**DOI:** 10.1101/2020.10.28.360065

**Authors:** Megha Murali, Sara R. Turner, Darrel Belke, William C. Cole, Justin A MacDonald

## Abstract

**Aims:** Smoothelin-like 1 (SMTNL1), a protein kinase A/G target protein, modulates the activity and expression of myosin light chain phosphatase and thus plays an important role in regulating vasoconstriction. Increased myogenic reactivity of resistance arterioles is associated with SMTNL1 silencing, and elevated baseline vascular tone is increasingly recognized as a risk factor for development of hypertension and chronic congestive heart failure. Hence, in this study we assessed cardiac function in SMTNL1 knockout mice with and without accompanying acute cardiac stress (i.e., pressure overload by transverse aortic constriction).

**Methods and Results:** Male and female, global *Smtnl1* knockout (KO) & wild-type (WT) mice were assessed at 10 weeks of age by echocardiography and electrocardiography to define baseline cardiac function. Gross dissection revealed distinct cardiac morphology only in male mice; hearts from KO animals were significantly smaller than WT littermates but the proportion of heart mass taken up by LV was greater. Non-invasive analyses of KO mice showed reduced resting heart rate with improved ejection fraction and fractional shortening as well as elevated aortic and pulmonary flow velocities relative to their WT counterparts, but only in the male cohort. We further investigated the impact of acute pressure overload on cardiac morphometry and hemodynamics in the absence of SMTNL1 in male cohort using echocardiography and pressure-volume (PV) loop measurements. Interestingly, PV loop analysis revealed diastolic dysfunction with significantly increased end diastolic pressure and LV relaxation time along with a steeper end diastolic pressure-volume relationship an indicator of stiffer heart, in the KO group when compared to WT Sham-operated group. Sham KO mice also showed elevated arterial elastance and total peripheral resistance. With acute pressure overload, systolic function was preserved, but diastolic dysfunction was exacerbated in KO mice with higher E/E’ ratio and myocardial performance index along with a prolonged isovolumetric relaxation time relative to the aortic-banded WT group.

**Conclusion:** Taken together, the findings support a novel, sex-dimorphic role for SMTNL1 in modulating cardiac structure and diastolic function. Significantly, impairment of diastolic function following pressure overload in young animals lacking SMTNL1 is mainly driven by increased systemic vascular resistance, which mimics the clinical pathophysiology of heart failure with preserved ejection fraction (HFpEF).

**Translational Perspective:** Heart failure with preserved ejection fraction (HFpEF) is characterized by the impairment of diastolic function and accounts for half of all heart failure cases. Unfortunately, there is as yet no proven therapy available for these patients as the pathophysiology is complicated with the presence of multiple comorbidities, microvascular dysfunction and a lack of an ideal animal model. The phenotype of *Smtnl1* global deletion male mice exhibits intriguing similarities to HFpEF, with elevated microvascular resistance driving diastolic dysfunction and LV remodeling. As such the SMTNL1 KO mouse represents a novel pre-clinical model to study the molecular etiology of HFpEF.

## 1. Introduction

Smoothelin-like 1 (SMTNL1; originally termed calponin homology-associated smooth muscle protein, CHASM) was first identified as a target of cGMP-dependent protein kinase (PKG) during Ca^2+^-desensitization of smooth muscle (Borman et al., 2004). However, unlike other smoothelin proteins which display restricted expression in smooth muscle cells, SMTNL1 can be found in other cell types, with the most abundant expression found in skeletal muscle (Wooldridge et al., 2008). Unique from other smoothelin proteins, SMTNL1 is phosphorylated by cyclic nucleotide-dependent protein kinases, PKA and PKG (Borman et al., 2004; Wooldridge et al., 2008). Maximal relaxation of aortic smooth muscle in response to acetylcholine was associated with phosphorylation of SMTNL1 at Ser301, an observation that alludes to an important role for SMTNL1 in the regulation of arterial tone (Wooldridge et al., 2008).

With the generation of a global *Smtnl1* knockout (KO) mouse, the physiological impact of SMTNL1 on the cardiovascular system was assessed (Wooldridge et al., 2008). SMTNL1 KO mice develop sex-dependent physiological adaptations to exercise and pregnancy in vascular, uterine and skeletal muscle. These phenotypes were attributed to the sex-dimorphic expression of SMTNL1, which increases throughout sexual development, with stable differences in expression levels observed in male and female mice regardless of age (Bodoor et al., 2011). Interestingly, SMTNL1 levels were also augmented in muscle tissues of wild-type female mice during pregnancy when compared to non-pregnant animals (Bodoor et al., 2011). Estrogen receptor (ER)-binding sites exist in the functional promoter region of the mouse *Smtnl1* gene, so sex hormones may also influence its transcriptional activity (Ulke-Lemée et al., 2011). Additionally, a substantial reduction in force development in response to contractile agonist phenylephrine, as well as sensitization to the relaxant agonist acetylcholine, was observed for vascular smooth muscle of SMTNL1 KO mice during pregnancy (Lontay et al., 2010).

The deletion of SMTNL1 in male mice also mimicked exercise-associated adaptations to the vasculature. Aortic rings from sedentary KO animals achieved the same extent of enhanced vasodilatory response to acetylcholine (~25% increase) as that from exercised wild-type animals (Wooldridge et al., 2008). Moreover, only male KO mice exhibited increased time to fatigue and more willingness to complete the treadmill running task with fewer motivational stimuli during the 5-week protocol, resembling a cardiovascular phenotype achieved from endurance-exercise training in a sex-dependent manner. It is noteworthy that exercise was found to reduce the expression of *Smtnl1* in wild-type mice (Wooldridge et al., 2008).

On the other hand, pressure myography experiments performed *ex vivo* on isolated resistance vessels such as cerebral and mesenteric arteries found vessels from male but not female KO mice to display enhanced myogenic reactivity in response to increasing luminal pressure (10-120 mmHg) while vessels from wild-type mice developed normal autoregulatory constrictions (Turner and MacDonald, 2014; Turner et al., 2019). No overt structural alterations were observed for vessels isolated from KO mice, and augmented PKC signaling acting on Ca^2+^-sensitizing mechanisms of smooth muscle contraction (i.e., myosin light chain phosphatase (MLCP) and the C-kinase potentiated inhibitor of myosin phosphatase (CPI-17)) were associated with enhanced myogenic reactivity of vessels in the absence of SMTNL1 (Turner et al., 2019). This finding was consistent with previous reports that revealed SMTNL1 to regulate myosin phosphatase activity through direct interaction with the myosin phosphatase-targeting subunit 1 (MYPT1) of MLCP (Borman et al., 2009; Lontay et al., 2010) and through transcriptional/translational effects on the expression levels of MYPT1 (Lontay et al., 2010).

While previous studies demonstrate substantive changes in vascular contractile performance in the absence of SMTNL1, none have specifically interrogated the impact of SMTNL1 on cardiac function. Hence, the goal of this study was to assess the effect of SMTNL1 silencing on cardiac morphometry and hemodynamics in both male and female cohorts. Furthermore, resistance vessels are critical in the establishment and maintenance of blood pressure (Burton, 1954; Harper and Chandler, 2016); enhanced myogenic tone of vessels from SMTNL1 KO mice is suggestive of increased total peripheral resistance, which could lead to high blood pressure. For this reason, we also examined the effect of acute pressure-overload driven by transverse aortic constriction (TAC) on cardiac function and morphology in the absence of SMTNL1. Herein, we report that male *Smtnl1*^-/-^ mice exhibit improved systolic function and altered cardiac morphometry compared to age-matched wild-type animals. A significantly reduced resting heart rate, altered electrical conduction properties along with increased pulmonary and aortic flow peak velocities in male SMTNL1 KO mice also imply that SMTNL1 influences cardiac hemodynamic performance in a sex-dimorphic way. Hemodynamic measurements revealed enhanced total peripheral resistance and mean arterial pressure in Sham-operated KO animals. Although the KO mice maintained a normal systolic function upon pressure overload associated with TAC, diastolic function was markedly impaired with evidence of enhanced concentric hypertrophy relative to wild-type mice; a phenotype consistent with the clinical condition of heart failure with preserved ejection fraction (HFpEF).

## 2. Methods

### 2.1 Animals

Wild-type (WT) and *Smtnl1* knockout mice (KO, global: previously described (Wooldridge et al., 2008) on a 129S6/SvEvTac background were employed. Mice were maintained on a normal chow diet (5062 Pico-Vac^®^ Mouse Diet 20) and standard 12L:12D light-dark cycle. Male and female mice were used at 10 weeks of age. All animal protocols and use were approved by the Animal Care and Use Committee at the University of Calgary and conform to the guidelines set by the Canadian Council of Animal Care. The investigators were blinded to genotype and sex of the mice for all data acquisition protocols and analyses.

### 2.2 Transverse aortic constriction

Male mice were anesthetized using isoflurane in oxygen (4% for induction; 2% for maintenance) and placed in a supine position on a heated platform to maintain normothermia. Analgesia was provided with subcutaneous injection of buprenorphine (0.05 mg/kg). TAC was achieved by tightening a suture against a blunted 27-gauge needle placed against the aorta (Tarnavski et al., 2004). For sham controls, the aorta was exposed in a cohort of mice but not banded. The mice were allowed to recover for 4-weeks prior to examination.

### 2.3 Echocardiography

Transthoracic echocardiography (Vevo770 & Vevo3100 instruments; Visual Sonics, Toronto, ON, Canada) was completed on mice maintained under general anesthesia. The mice were depilated as required for imaging and placed on a heated platform to maintain normothermia for the duration of the procedure. M-Mode imaging was used to obtain measurements of left ventricle posterior wall thickness (LVPWT), interventricular septal thickness (IVS) as well as LV internal dimensions (LVID) in diastole and systole. Fractional shortening (FS%) and ejection fraction (EF%) were calculated by averaging values obtained from 5 cardiac cycles. Pulsed-wave Doppler was used in long-axis imaging mode to obtain mean flow velocities and velocity-time integrals for the ascending aortic arch as well as pulmonary artery. Diastolic function was measured by conventional Doppler analysis of peak early- and late-diastolic transmitral velocities (Gao et al., 2011). All measurements were made from original tracings and 5-10 beats were averaged for each measurement.

### 2.4 Electrocardiography

Recordings were obtained by lightly restraining the mice in a small electrocardiography (ECG) tube apparatus (QRS Phenotyping Inc., Calgary). Mice were acclimated to the apparatus for 15 min, and then ECGs were acquired with to define heart rate (HR), R-R, P-R, QRS, J-T(c), Q-T(c) and TpTe intervals, and amplitudes. The data were manually checked for outliers due to animal movement.

### 2.5 Hemodynamics and assessment of pressure-volume relationships

Mice were secured on a heated pad to maintain normothermic body temperature, and the LV was catheterized through the right carotid artery to simultaneously measure the pressure and the volume, using an ultraminiature 1 F impedance-pressure catheter (PVR 1045, Millar Instruments, Houston, Texas). Closed-chest pressure-volume (PV) recordings were obtained at steady state, with and without ventilation, and during transient occlusion of the inferior vena cava to obtain preload-independent indexes (Pacher et al., 2008). The PV data were collected at a sampling rate of 1 kHz using an MPVS Ultra system (Millar Instruments, Houston, Texas) in tandem with a Power Lab data acquisition system (AD Instruments Inc., Colorado, USA). Mean arterial pressure (MAP) and systolic pressure (SP) was recorded from the aorta at the start of the experiment, and the total peripheral resistance (TPR) [MAP/cardiac output] was calculated.

### 2.6 Histology

Hearts were excised, perfused with 10% buffered formalin, and immersion fixed for a week at room temperature prior to paraffin embedding. Serial sections (5 μm thick) were stained with hematoxylin and eosin (H&E) and picrosirius red (PSR). The area of collagen deposition, excluding the perivascular fibrotic area was quantified using ImageJ, by averaging measurements from 3 different regions of each mouse heart (i.e., intraventricular septal wall, LV posterior wall and apex).

### 2.7 Serum biomarkers

Blood was collected by cardiac puncture immediately following euthanasia. Serum was separated using standard methods and stored as aliquots at −80 °C. A Mouse Cardio 7-plex (Eve Technologies, Calgary, AB, Canada) assay was carried out to quantify systemic cardiovascular inflammatory biomarkers.

### 2.8 PCR and western immunoblotting

Total RNA was extracted and purified from isolated cardiac muscle and gastrocnemius skeletal muscle tissues. The relative expression levels of *Smtnl1* were normalized against 18S-rRNA. Each sample was measured in triplicate. For western blots, clarified tissue homogenates were prepared for analyses. Samples were subjected to SDS-PAGE (10%) and then transferred to nitrocellulose (0.2 μm). The membranes were blocked with 5% w/v non-fat milk and then incubated with primary antibody (pAb #1, rabbit anti-SMTNL1 IgG raised against a ^206^DTKELVEPESPTEEQ^220^ peptide, 1:1000; pAb #2, rabbit anti-SMTNL1 IgG raised against mouse SMTNL1^1-346^ protein lacking the calponin homology (CH)-domain, 1:500). The blots were washed extensively prior to incubation with HRP-conjugated secondary antibody (1:2000). The blots were developed with enhanced chemiluminescence, and densitometry was completed with an LAS4000 Imager and ImageQuantTL software (GE Healthcare Inc).

### 2.9 Statistical Analysis

Data are presented as mean ± SEM, and the statistical analyses were performed with GraphPad Prism 8.3.0 software. The Student’s unpaired t-test was used to compare between 2 groups, and ANOVA (two-way) followed by Tukey’s multiple comparison test was used to assess more than 2 groups.

## 3. Results

### 3.1 Male *Smtnl1^-/-^* mice have smaller cardiac dimensions but improved systolic function

In order to investigate the potential cardiac manifestations of SMTNL1 silencing, we evaluated heart morphology and cardiac function in both male and female, wild-type (WT) and SMTNL1 knockout (KO) mice at 10 weeks of age. A marked decrease in gross heart dimension and mass (**Figure 1A** and **1B**, respectively) and significant decreases in heart mass (HM) to body mass (BM), and HM to tibia length (TL) ratios (**Figure 1C**) were detected between WT and KO male mice. Conversely, the left ventricular mass (LVM) as a proportion of the total HM was found to be elevated in hearts of male KO mice (**Figure 1D**). In all cases, female hearts were found to be significantly smaller than those of males; however, no differences in gross heart dimensions, HM:BM, HM:TL or LVM/HM, were observed with knockout of SMTNL1 in young female mice. *Smtnl1* expression was low in heart tissue with transcript levels ~500-fold less than that found in skeletal muscle (**Figure 1E**). SMTNL1 protein could not be identified in LV tissue lysates at the expected molecular weight using antibodies raised against two different epitopes (**Figure 1F**). Furthermore, the levels of serum markers of vasculopathy (i.e., sE-selectin and sP-selectin; elevated in KOs), cardiac remodeling (i.e., pro-matrix metalloprotease 9 (pro-MMP9); elevated in KOs) and angiogenesis (i.e., vascular endothelial growth factor (VEGF); decreased in KOs) were significantly impacted by global deletion of SMTNL1 (**Figure 1G**).

**Figure 1:**
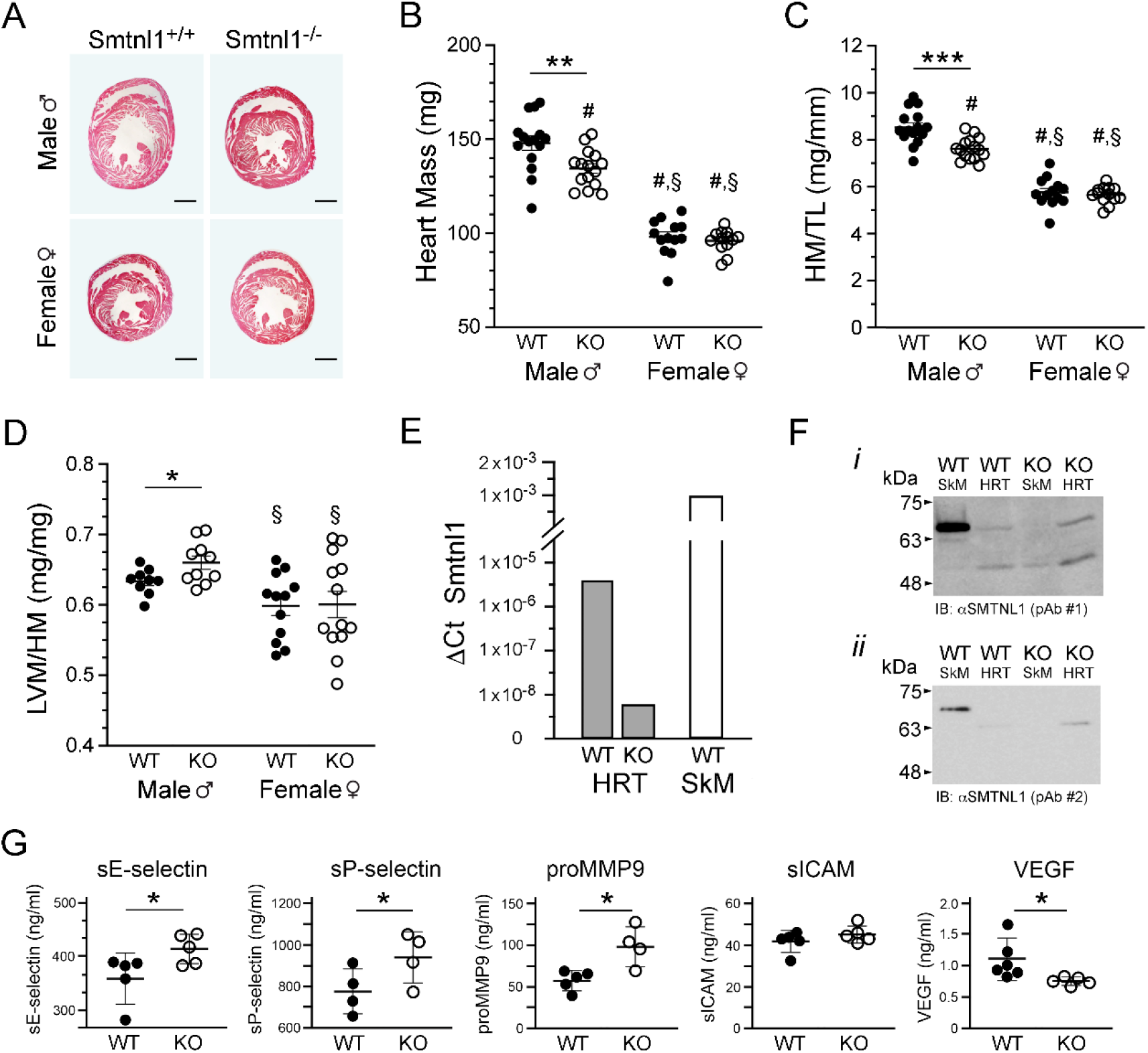
Morphometric cardiac analysis of male and female SMTNL1 knockout (KO) mice. Representative H&E-stained transverse sections of hearts dissected from *Smtnl1*^-/-^ (KO) and *Smtnl1*^+/+^ (WT) mice (**A**); scale bar, 1 mm. Male and female mice were fed a normal diet, held in separate cages after weaning, and studied at 3 months of age. Gross heart mass (HM; **B**) as well as ratios of heart mass to tibia length (HM:TL, **C**) and left ventricular mass to heart mass (LVM:HM, **D**) are provided. The data represent means ± SEM, n=13-15 mice per experimental group. In (**E**), qPCR analysis of Smtnl1 gene expression in cardiac and skeletal muscle tissues of male WT and KO mice. Relative quantification (ΔCt) of *Smtnl1* expression was completed using 18S-rRNA as the normalization standard. In (**F**), western blot analysis of isolated cardiac muscle (HRT, 65 μg total protein load) or skeletal muscle (SkM, 25 μg total protein load) lysates obtained from male WT and KO mice. Two different primary polyclonal antibodies generated against mouse SMTNL1 were used: (*i*), rabbit anti-SMTNL1 IgG raised against the ^206^DTKELVEPESPTEEQ^220^ peptide, and (*ii*) rabbit anti-SMTNL1 IgG raised against SMTNL1^1-346^ mouse protein lacking the calponin homology (CH)-domain. The images are representative of two independent experimental assessments. In (**G**), the levels of circulating biomarkers were quantified by multiplexed LASER bead assay of serum obtained from male WT and KO mice (n = 4-6). Significantly different from WT males (#) or KO males (§) by two-way ANOVA with Tukey *post hoc* analysis (P < 0.01). Significantly different from WT males by two-tailed Student’s t-test: * – P < 0.05, ** – P < 0.01, and *** – P < 0.001.

We performed 2D M-mode echocardiography to examine the cardiac structure and function of *Smtnl1*^-/-^ and WT mice (**Figure 2A**). Significant differences in parameters associated with the pumping function of the heart and its shape were identified by Echo. Interestingly, increases in LV fractional shortening (FS%, **Figure 2B**) and LV ejection fraction (EF%, **Figure 2C**) were observed for male *Smtnl1*^-/-^ mice when compared to WT groups. No impact of SMTNL1 silencing was observed for cardiac function parameters when WT and KO female mice were compared. The LV end-diastolic diameter (LVIDd) and LV end-systolic diameter (LVIDs) were both smaller in the male KO mice when compared with the male WT group (**Table 1**). Moreover, LV wall thicknesses were consistently larger in KO male mice, with the end-systolic interventricular septum (IVSs) thickness reaching statistical significance when compared to the WT. Although LV diameters of WT female animals were smaller than corresponding WT males, no differences in cardiac structure (e.g., LVID, LVPW, and IVS) were identified between WT and KO animals.

**Figure 2:**
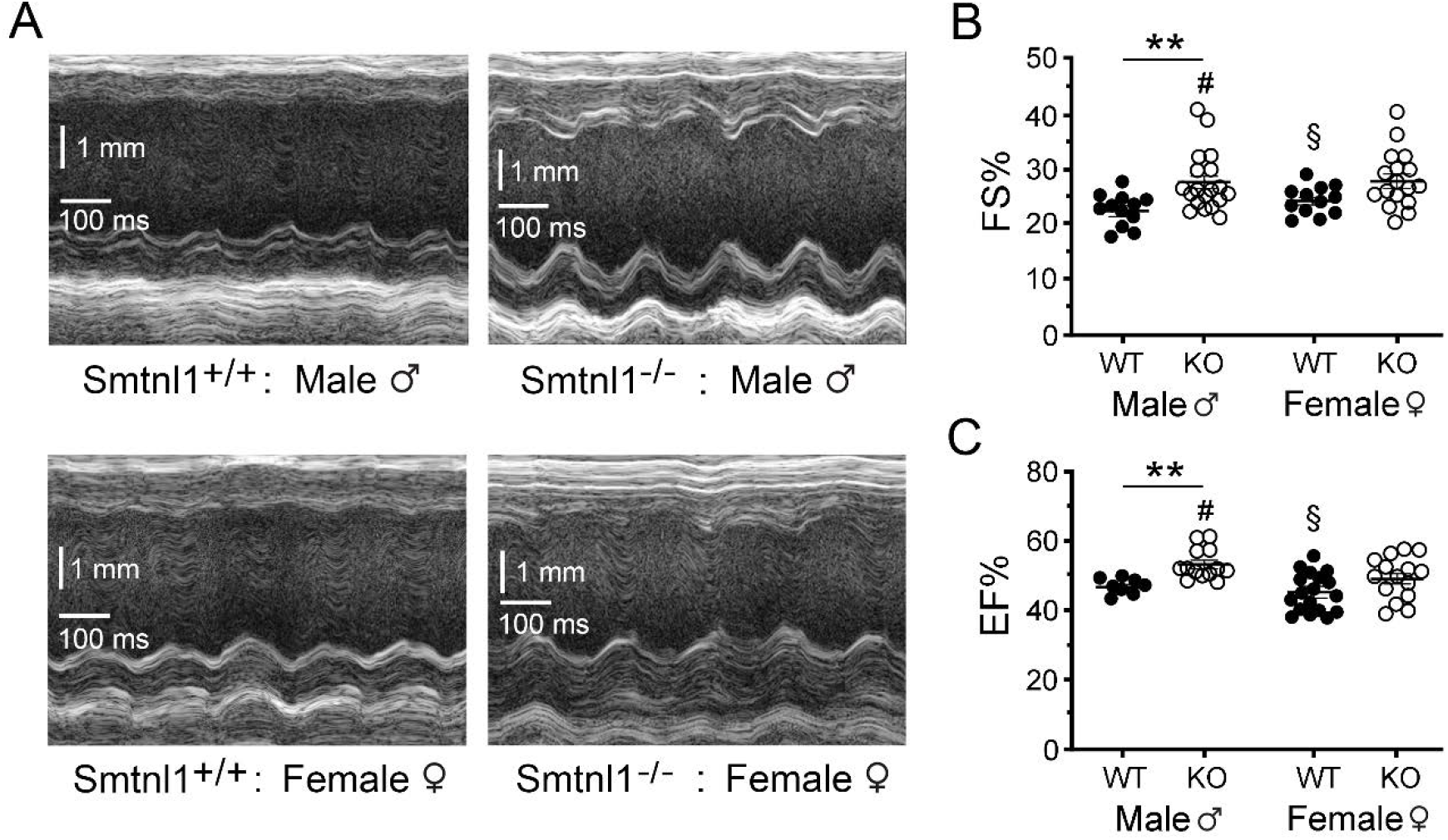
Echocardiographic analysis of ventricular morphology in SMTNL1 knockout (KO) mice. Representative M-mode echocardiograms **(A)** collected from both male and female, WT and KO mice at 3 months of age. Cumulative data of fractional shortening (FS%) and LV ejection fraction (EF%) are shown in panels **(B)** and **(C)**, respectively. The data represent means ± SEM, n=8-19 mice per experimental group. Significantly different from WT males (#) or KO males (§) by two-way ANOVA with Tukey *post hoc* analysis (P < 0.01). Significantly different from WT males by two-tailed Student’s t-test: ** – P < 0.01.

**Table 1.** Echocardiography of wild-type (WT) and *Smtnl1*^-/-^ (KO) hearts.

|  | Male ♂ |  | Female ♀ |  |
| --- | --- | --- | --- | --- |
|  | *Smtnl1*<sup>+/+</sup> (WT) | *Smtnl1*<sup>-/-</sup> (KO) | *Smtnl1*<sup>+/+</sup> (WT) | *Smtnl1*<sup>-/-</sup> (KO) |
| FS% | 22.5 ± 0.8 | 27.5 ± 1.2* | 24.1 ± 0.8 | 26.9 ± 1.1 |
| EF% | 47.2 ± 0.7 | 53.4 ± 1.2 * | 46.4 ± 1.3 | 49.5 ± 1.6 |
| IVSd | 0.80 ± 0.04 | 0.93 ± 0.06 | 0.89 ± 0.06 | 0.88 ± 0.04 |
| IVSs | 1.09 ± 0.05 | 1.29 ± 0.05* | 1.15 ± 1.15 | 1.17 ± 0.04 |
| LVPWd | 0.82 ± 0.02 | 0.95 ± 0.07 | 0.98 ± 0.98 | 1.00 ± 0.07 |
| LVPWs | 1.08 ± 0.03 | 1.22 ± 0.06* | 1.23 ± 0.10 | 1.28 ± 0.07 |
| LVIDd | 4.24 ± 0.707 | 3.98 ± 0.10 | 3.86 ± 0.09 | 3.82 ± 0.07 |
| LVIDs | 3.41 ± 0.08 | 2.98 ± 0.09* | 3.09 ± 0.07 | 2.95 ± 0.07 |

The left ventricles of WT and SMTNL1 KO mice were examined using M-mode echocardiography. FS: fractional shortening; EF: ejection fraction; IVSd, IVSs: interventricular septum during diastole or systole, respectively; LVPWd, LVPWs: left ventricular posterior wall during diastole or systole, respectively; LVIDd, LVIDs: left ventricular internal diameter during diastole or systole, respectively. Values are expressed as means ± SEM, n = 8-19 mice per experimental group. Significantly different from WT group: \*- P < 0.05, two-tailed Student's t-test.

### 3.2 Male *Smtnl1*^-/-^ mice display increased aortic and pulmonary blood flow

Blood flow through the ascending aortic arch was assessed using pulsed-wave Doppler flow analysis (**Figure 3A)**. Peak aortic arch velocity (AoAV) increased from 1481 ± 94 mm/s in the WT male to 1898 ± 95 mm/s in the KO male (**Figure 3A, ii**). Mean AoAV was also increased by ~250 mm/s in the KO male when compared to WT males. Increases in the peak aortic flow pressure gradient (AoPG, **Figure 3A, iii**) of ~6 mmHg and the velocity–time integral (VTI, **Figure 3A, iv**) of ~ 2.0 cm/systole were also observed. Data were analyzed by two-way ANOVA, which was negative for an interaction, genotype and biological sex factors (p > 0.05); however, Student’s *t*-tests performed against WT and KO males showed significant increases in AoAV values (peak, p = 0.027; mean, p = 0.028), AoPG values (peak, p = 0.042; mean, p = 0.044) and AVI (p = 0.045). No differences were observed between WT and KO female mice. Pulmonary arterial flow was assessed in the same manner, and representative recordings are presented in **Figure 3B**. Peak pulmonary arterial flow velocity (PAV, **Figure 3B, ii**) was increased with SMTNL1 knockout in male mice (i.e., WT, 632 ± 34 mm/s; KO, 731 ± 29 mm/s; Student’s t-test, p = 0.045). Peak pulmonary flow pressure gradient (PAPG, **Figure 3B, iii**) was also increased in male SMTNL1 knockout mice (i.e., WT, 1.64 ± 0.15 mmHg; KO, 2.17 ± 0.17 mmHg; Student’s t-test, p = 0.036); however, the VTI was not affected by *Smtnl1* deletion in this case (**Figure 3B, iv**). No differences were observed between WT and KO female mice. Data were analyzed with two-way ANOVA, and there was not a significant interaction between genotype and sex. However, the biological sex factor was significant (p = 0.035) in that female animals had lower peak pulmonary flow regardless of genotype. Moreover, the two-way ANOVA was also significant for the genotype factor (p = 0.030) with KO animals demonstrating higher pulmonary arterial flow regardless of sex.

**Figure 3:**
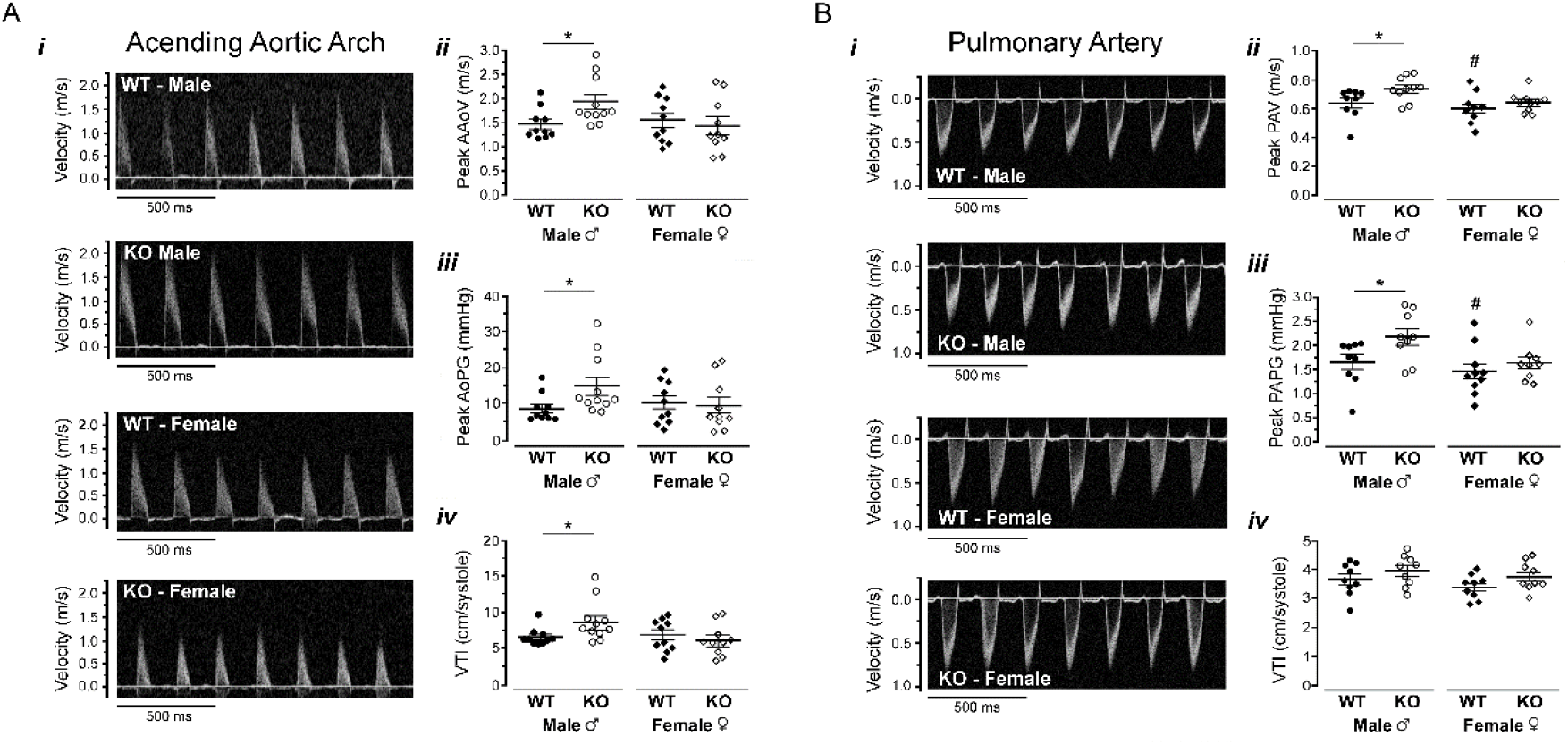
Evaluation of blood flow velocity of the ascending aorta and pulmonary artery by Doppler imaging. Representative pulse-wave Doppler echocardiograms for the ascending aortic arch **(A, i)** and pulmonary artery (**B, i)** were collected for male and female, WT and KO mice at 10 weeks of age. Cumulative data are provided for peak ascending aortic arch flow velocity (AAoV; **A, ii**), peak aortic pressure gradient (AoPG; **A, iii**), the velocity-time integral (VTI,; **A, iv**), peak pulmonary artery flow velocity (PAPV; **B, ii**), peak pulmonary artery pressure gradient (PAPG; **B, iii**), and the velocity-time integral (VTI; **B, iv**). * - indicates a significant difference between WT and KO male animals by two-tailed Student’s t-test (P < 0.05; n=10). # - indicates significantly different from male KO mice by two-way ANOVA with Tukey post-hoc analysis (P < 0.05, n=10).

### 3.3 *Smtnl1*^-/-^ is associated with variations in cardiac electrophysiological parameters

To further investigate the effect of global SMTNL1 deletion on cardiac function, electrocardiograms (ECGs) were performed. Cumulative data for heart rate and wave intervals of 10-week-old, male, WT and KO animals (*n* = 12) are presented in **Figure 4**. Since SMTNL1 deletion did not provide distinction in cardiac form or function in female animals, ECGs were not completed on this cohort. Heart rate was significantly lower in KO males, 625.3 ± 22.6 beats/min compared with 687.9 ± 6.9 beats/min (**Figure 4A**; p = 0.015) and, as expected, was accompanied with an increased R to R interval between heart beats (**Figure 4B**; p = 0.028). Interestingly, while the interval between heartbeats was lengthened, atrial depolarization of the heart appeared to be faster in KO animals as evidenced by the decreased duration of P waves (**Figure 4C**, p = 0.011). The interval from the peak of the T wave to the end of the T wave (Tpeak to Tend; TpTe) provides an approximation of transmural dispersion of ventricular repolarization (Antzelevitch, 2007); thus, ventricle repolarization also appeared more rapid in male SMTNL1 KO mice with diminished TpTe intervals (**Figure 4D**; p = 0.018). There was no change in the PR interval (**Figure 4E**), so wave propagation through the atrioventricular node appeared unaffected by SMTNL1 deletion. The QRS complex interval (**Figure 4D**) was also unchanged, suggesting no impact of SMTNL1 deletion on ventricular depolarization rates. A shortened QT interval was also identified in male *Smtnl1*^-/-^ mice (**Figure 4G**; p = 0.041), representing a decrease in the period of ventricular systole from ventricular isovolumetric contraction to isovolumetric relaxation (Speerschneider et al., 2013; Roussel et al., 2016). Regardless, SMTNL1 deletion was also associated with shortened QTc intervals after correcting for differences in the heart rates (data not shown). Likewise, the JT interval (**Figure 4H**; and JTc when corrected for heart rate) was also shortened in the SMTNL1 KO mice, suggesting faster ventricular repolarization. Finally, the waveform amplitudes were also analyzed, and summative data are provided (**Figure 4I-L)**. Only the S wave, of the QRS complex, was significantly impacted by SMTNL1 deletion, with KO animals having a greater waveform magnitude (**Figure 4I**; p = 0.010). In summary, the results of ECG analyses suggest that male SMTNL1 KO animals have longer intervals between heart beats, but faster propagation of electrical signals through the atria and ventricles (shortened P and T waves), with minimal to no changes in the waveform amplitudes.

**Figure 4:**
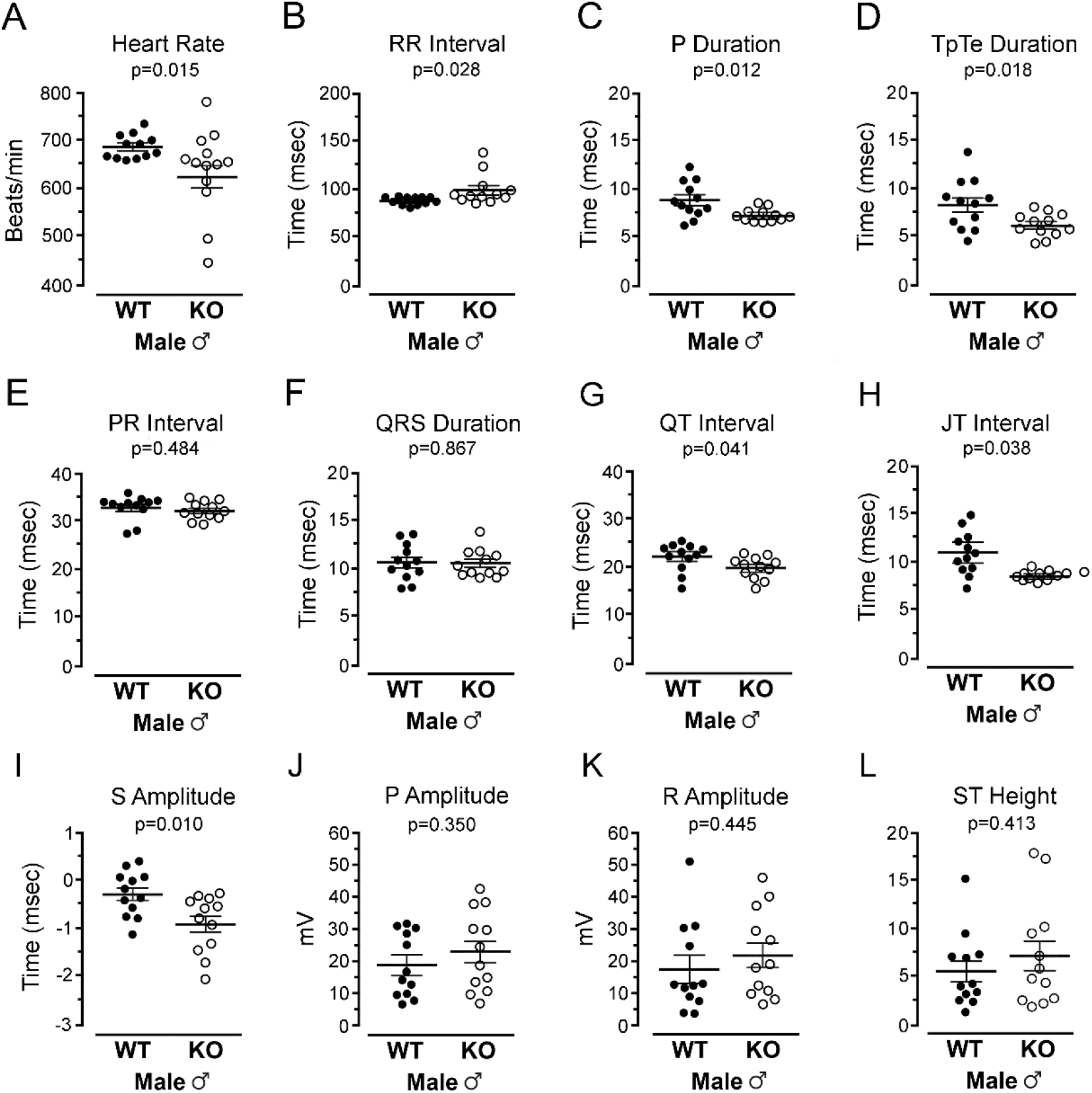
Impact of SMTNL1 deletion on cardiac electrophysiological parameters. A surface electrocardiogram (ECG) was used to monitor heart rate (**A**) and cardiac wave form properties in conscious male, 10-week-old WT and KO mice. Various ECG parameters are indicated: R-R interval (**B**), P duration (**C**), Tpeak to Tend (TpTe) duration (**D**), PR interval (**E**), QRS duration (**F**), as well as QT and JT intervals (**G** and **H**), The amplitude of each cardiac wave form was also measured (mV). The S wave amplitude (**I**) was significantly enhanced in KO male animals, but no differences in P amplitude (**J**), R amplitude (**K**) or ST height (**L**) were observed with SMTNL1 deletion. Data were analyzed by Student’s *t-* test (n = 12); P values are indicated on graphs with P < 0.05 indicating statistical significance.

### 3.4 Pressure overload elicits increased cardiac hypertrophy in *Smtnl1*^-/-^ mice

Following the baseline characterization of the SMTNL1 KO mice, we hypothesized that these mice would be more susceptible to cardiac stress elicited with a pressure-overload model of left-sided heart failure (Tarnavski, 2009; Tarnavski et al., 2004). Hence, TAC was applied for 4 weeks to assess the impact of pressure overload on the early stages of hypertrophic remodeling. The mass of *Smtnl1*^-/-^ hearts was smaller than the WT in the sham groups (HW/BW, 5.1 ± 0.07 mg/g [KO] vs 5.6 ± 0.04 mg/g [WT], p = 0.012) (**Figure 5A**); a finding that was consistent with the baseline heart morphology (**Figure 1**). TAC elicited hypertrophic enlargement of KO hearts so that they became similar in size to WT hearts (**Figure 5A-C**). The KO mice possessed smaller hearts at the beginning of the 4-week TAC period, so the normalized hypertrophic growth was greater for the *Smtnl1*^-/-^ mice when compared to WTs. The provision of TAC also caused comparable increases in fibrotic deposition in the interstitial spaces of the LV myocardium for both WT (1.15 ± 0.15% [Sham] vs 20.14 ± 2.87% [TAC], p = 0.0001) and KO hearts (1.49 ± 0.23% [Sham] vs 15.73 ± 1.52% [TAC], p = 0.006) (**Figure 5D & 5E**).

**Figure 5.**
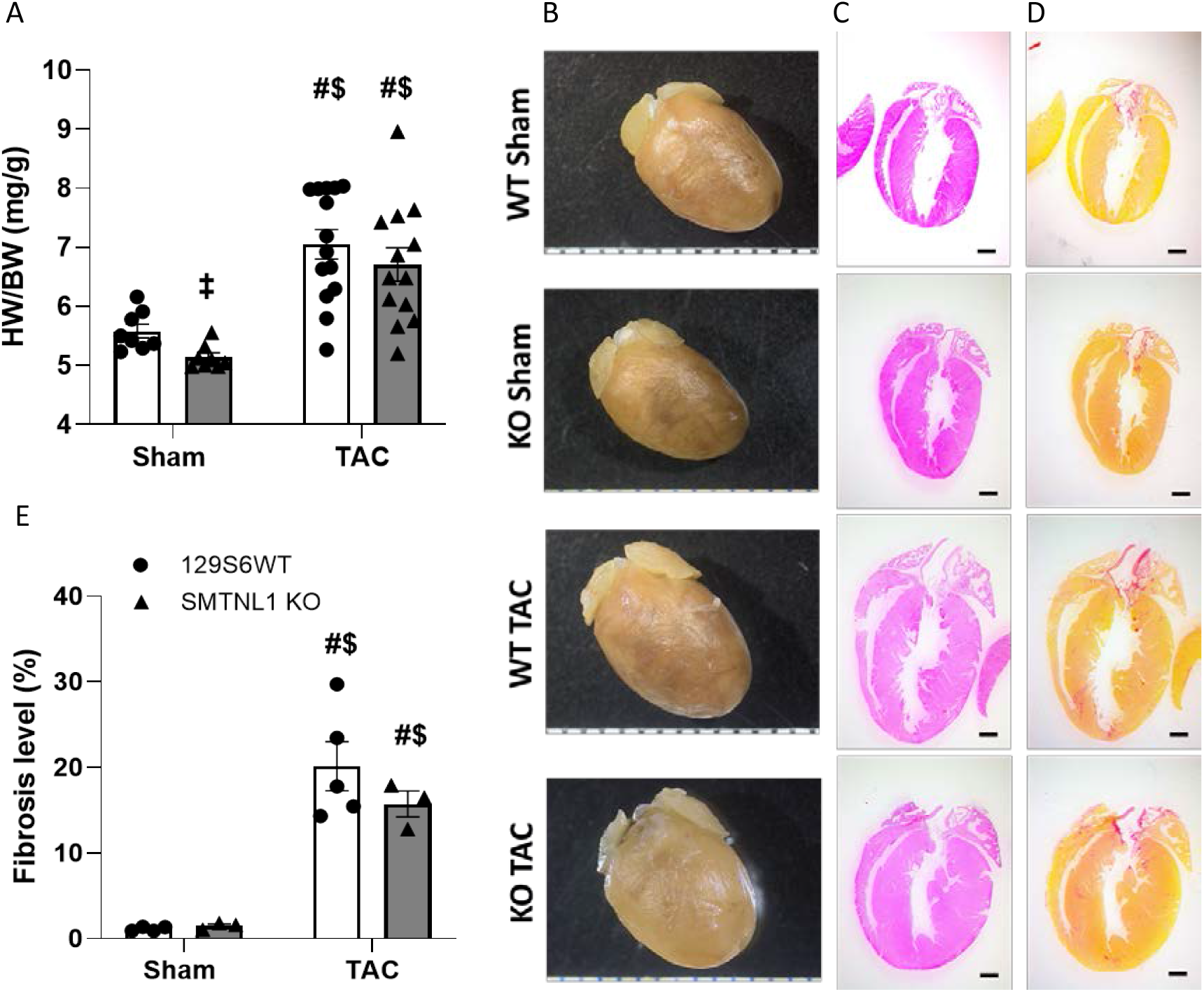
Impact of pressure overload on cardiac morphometry and fibrosis in male SMTNL1 knockout (KO) mice. (**A**) Heart weight to body weight ratio (n=7-8/group), Representative images of (**B**) gross morphology of whole hearts, (**C**) H&E and (**D**) picrosirius stained whole heart sections from pressure overloaded and sham controls, (**E**) quantified fibrotic levels (n=3-5/group). Scale bar, 1 mm. Data presented as mean ± SEM as analyzed by two-way ANOVA followed by Tukey’s multiple comparison test. ‡ - P < 0.05 vs WT control; # – P < 0.05 vs Sham WT; $ – P < 0.05 vs Sham KO.

### 3.5 *Smtnl1*^-/-^ mice exhibit cardiac remodeling with preserved systolic function following aortic constriction

Echocardiographic measurements corroborate the development of cardiac hypertrophy post-TAC in both genotypes with significantly enlarged IVS wall, LV posterior wall, LV relative wall thickness and LV mass when compared to their Sham counterparts (**Table 2**). The proportional change in LV relative wall thickness from the Sham counterpart was greater for the TAC KO cohort than for the TAC WT group (**Figure 6A**; 0.14 ± 0.03 mm [WT] vs 0.23 ± 0.03 mm [KO], p = 0.0384), and the decrease in the LVID during systole (**Figure 6B**; −0.035 ± 0.115 mm [WT] vs −0.312 ± 0.097 mm [KO], p = 0.0816) and diastole (**Figure 6C**; −0.051 ± 0.115 mm [WT] vs −0.338 ± 0.097 mm [KO], p = 0.0724) was more apparent in the TAC KO cohort. Taken together, the data support the selective development of concentric hypertrophy in hearts of *Smtnl1*^-/-^ animals after 4 weeks of TAC with no conspicuous changes in systolic functional parameters such as EF%, FS%, cardiac output, stroke volume, stroke work, preload recruited stroke work and the end systolic pressure volume relationship **(Table 2)**.

**Figure 6.**
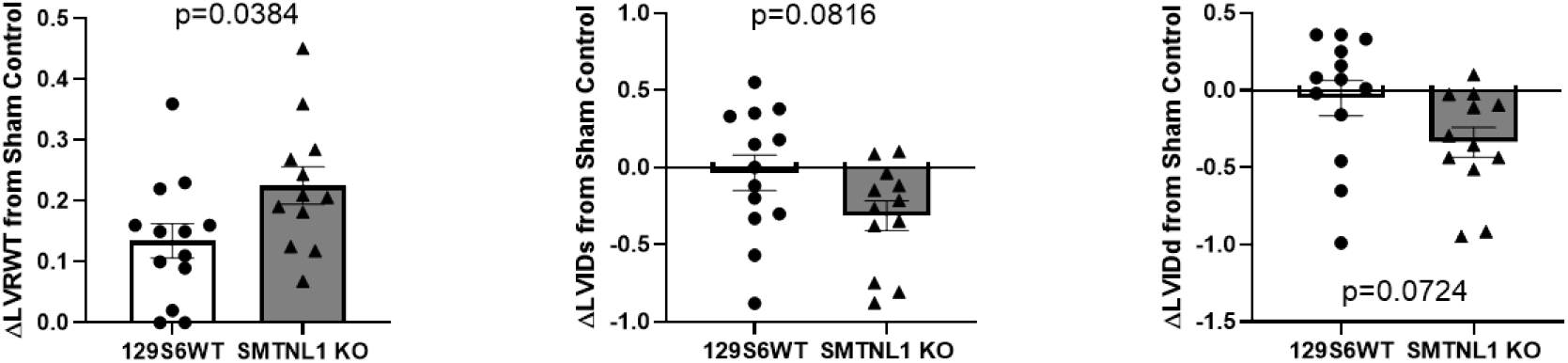
Change in cardiac structural parameters between TAC-banding and Sham controls for WT and KO SMTNL1 mice. (**A**) LVRWT, left ventricular relative wall thickness; (**B**) LVIDs, LV internal diameter during systole, and (**C**) LVIDd, LV internal diameter during diastole values were obtained with M-mode echocardiography. The differences (Δ) were calculated by subtracting the mean value determined for the WT and KO Sham group from the respective TAC-banded group values. Data were analyzed by Student’s *t*-test and presented as mean ± SEM, n=12-13/group. P values are indicated on graphs with P < 0.05 indicating statistical significance.

**Table 2.**
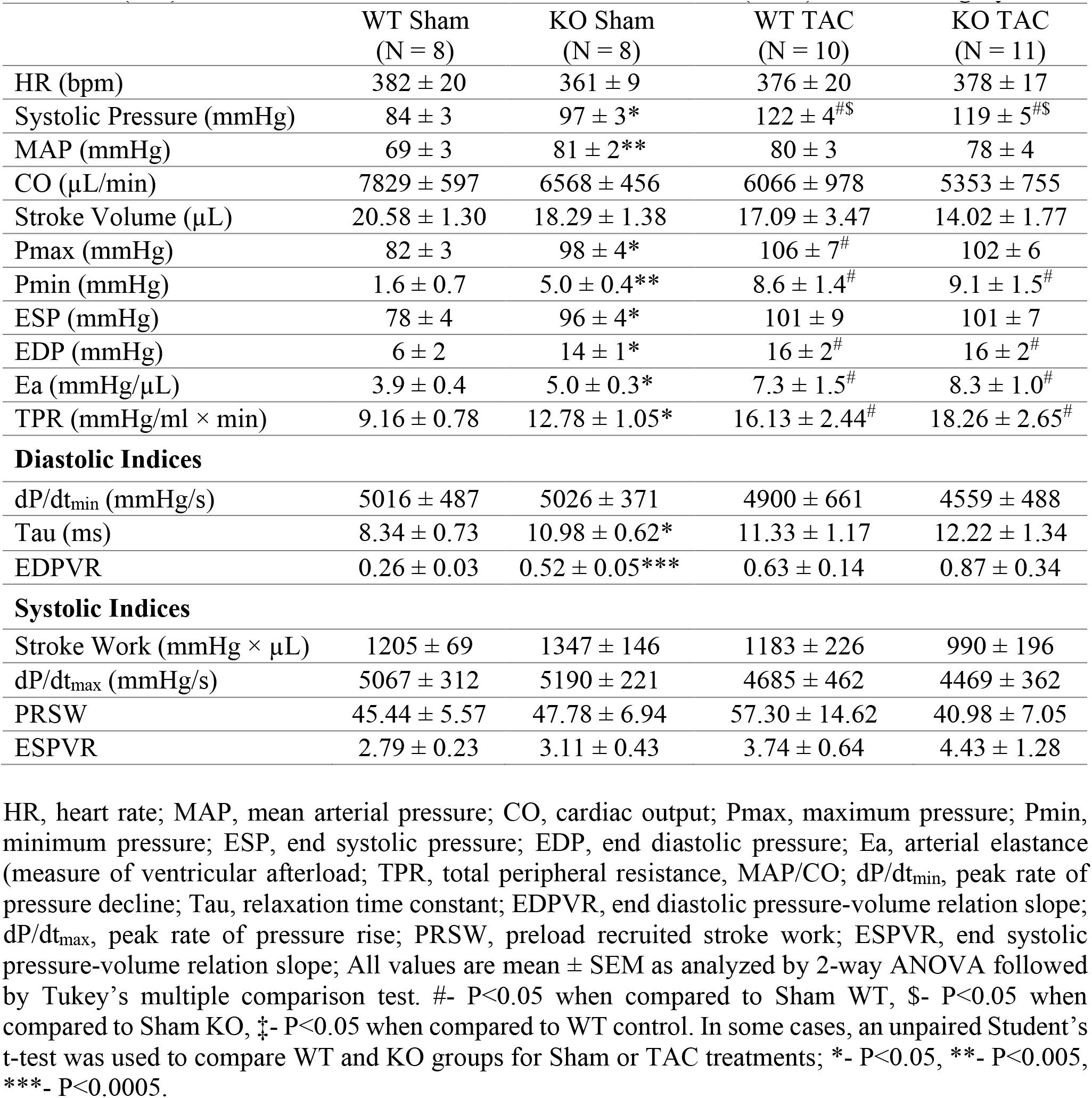
Hemodynamic parameters derived from PV relations for wild-type (WT) and *Smtnl1*^-/-^ (KO) mice 4 weeks after transverse aortic constriction (TAC) or sham surgery.

### 3.6 SMTNL1 silencing is associated with elevated pressures, increased arterial elastance and peripheral resistance

The systolic (SP, 97 ± 3 mmHg [KO] vs. 84 ± 3 mmHg [WT], p = 0.006) and mean arterial pressures (MAP, 81 ± 2 mmHg [KO] vs. 69 ± 3 mmHg [WT], p = 0.004) along with maximum (Pmax) and minimum pressures (Pmin) obtained from the PV loop measurements were significantly elevated in Sham KO when compared to Sham WT group (**Table 3**); however, the pressures measured after 4 weeks of TAC were comparable between the genotypes. Ultimately, a greater proportional increase in SP (**Figure 7A**; 38 ± 4 mmHg [WT] vs. 22 ± 5 mmHg [KO], p = 0.025), MAP (**Figure 7B**; 11 ± 3 mmHg [WT] vs. −3 ± 4 mmHg [KO], p = 0.011) and Pmax (**Figure 7C**; 24 ± 7 mmHg [WT] vs. 4 ± 6 mmHg [KO], p = 0.034) occurred for the TAC WT mice when the genotypes were compared with the corresponding Sham controls. Vascular indices such as arterial elastance (Ea, 5.0 ± 0.3 mmHg/μl [KO] vs. 3.9 ± 0.4 mmHg/μl [WT], p = 0.039) and total peripheral resistance (TPR, 12.78 ± 1.05 mmHg/ml × min [KO] vs. 9.16 ± 0.78 mmHg/ml × min [WT], p = 0.018) (**Table 2**) were found to be elevated for *Smtnl1*^-/-^-mice within the Sham cohorts, with a trend of further increase for both genotypes following 4 weeks of TAC. But there was no distinction identified between WT and KO cohorts for these values following TAC.

**Figure 7.**
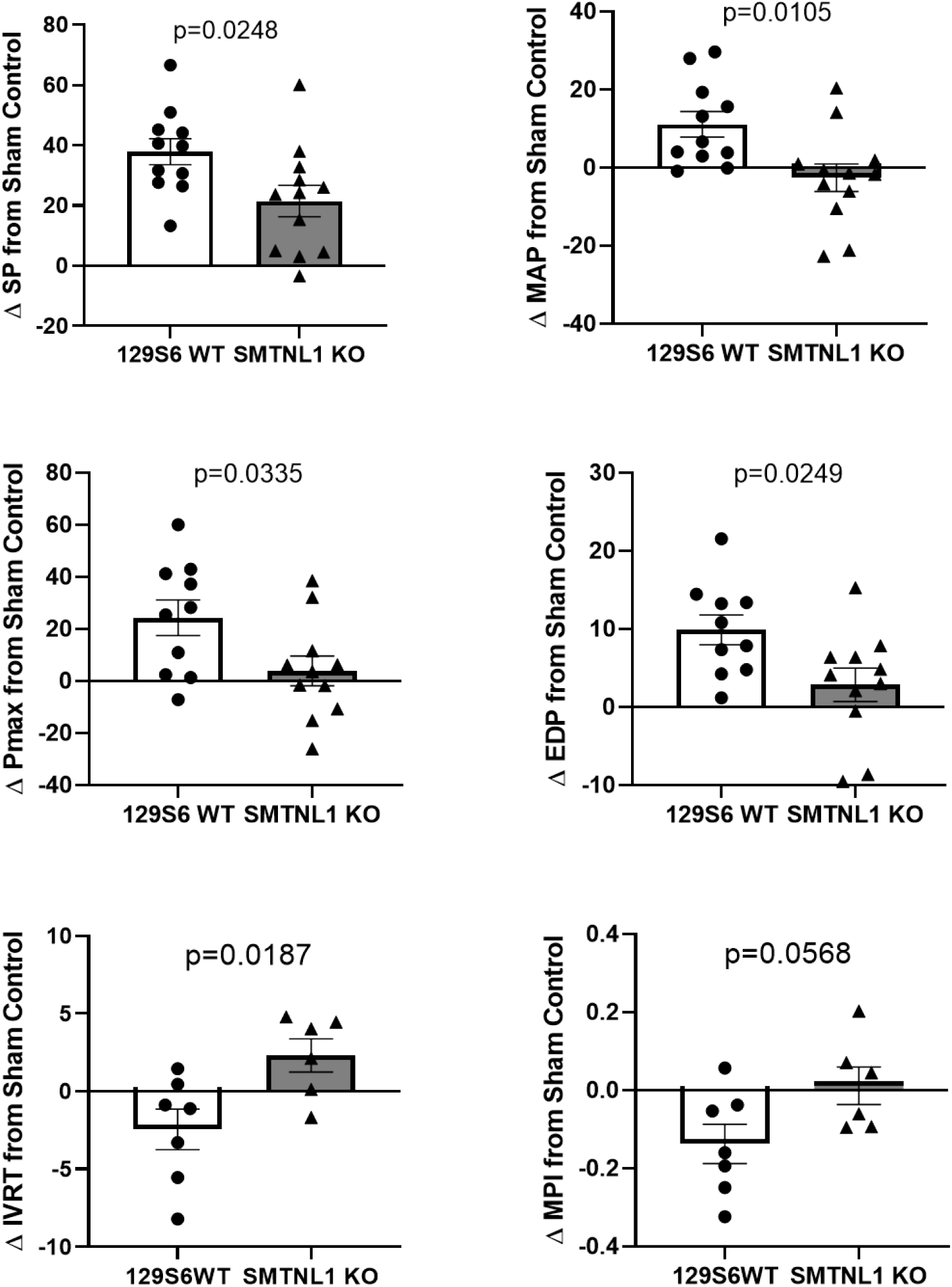
Change in cardiac hemodynamic parameters between TAC-banding and Sham controls for WT and KO SMTNL1 mice. (**A**) SP, systolic pressure; (**B**) MAP, mean arterial pressure; (**C**) Pmax, maximum pressure; (**D**) EDP, end diastolic pressure; (**E**) IVRT, isovolumetric relaxation time; and (**F**) MPI, myocardial performance index values were obtained from pressure-volume relationships of anesthetized mice. The differences (Δ) were calculated by subtracting the mean value determined for the WT and KO Sham group from the respective TAC-banded group values. Data were analyzed by Student’s *t*-test and presented as mean ± SEM, n=10-11/group. P values are indicated on graphs with P < 0.05 indicating statistical significance.

**Table 3.** Echocardiographic measurements for wild-type (WT) and *Smtnl1*^-/-^ (KO) mice 4 weeks after transverse aortic constriction (TAC) or sham surgery.

|  | WT Sham<br>(N = 8) | KO Sham<br>(N = 8) | WT TAC<br>(N = 10) | KO TAC<br>(N = 11) |
| --- | --- | --- | --- | --- |
| LVIDs (mm) | 3.20 ± 0.11 | 3.29 ± 0.12 | 3.16 ± 0.12 | 2.98 ± 0.10 |
| LVIDd (mm) | 4.38 ± 0.08 | 4.47 ± 0.12 | 4.33 ± 0.12 | 4.13 ± 0.10 |
| IVSs (mm) | 1.17 ± 0.09 | 1.17 ± 0.10 | 1.47 ± 0.10 | 1.57 ± 0.08<sup>#</sup> |
| IVSd (mm) | 0.73 ± 0.07 | 0.72 ± 0.08 | 1.02 ± 0.08<sup>#</sup> | 1.12 ± 0.07<sup>#</sup> |
| LVPWs (mm) | 1.23 ± 0.10 | 1.11 ± 0.02 | 1.37 ± 0.07 | 1.38 ± 0.09 |
| LVPWd (mm) | 0.79 ± 0.06 | 0.70 ± 0.03 | 1.06 ± 0.07<sup>#</sup> | 1.12 ± 0.05<sup>#</sup> |
| LVRWT | 0.35 ± 0.03 | 0.32 ± 0.03 | 0.48 ± 0.03<sup>#</sup> | 0.55 ± 0.03<sup>#</sup> |
| LV mass (mg) | 128.9 ± 11.5 | 120.2 ± 6.3 | 198.2 ± 19.8<sup>#</sup> | 198.8 ± 11.9<sup>#</sup> |
| EF% | 52.8 ± 2.3 | 51.8 ± 2.0 | 52.8 ± 1.9 | 54.4 ± 2.0 |
| FS% | 27.1 ± 1.4 | 26.5 ± 1.2 | 27.1 ± 1.1 | 28.0 ± 1.3 |
| IVCT (ms) | 19.3 ± 4.6 | 16.5 ± 2.1 | 17.9 ± 3.4 | 25.0 ± 1.4 |
| IVRT (ms) | 26.5 ± 1.2 | 26.5 ± 2.2 | 24.1 ± 1.3 | 28.9 ± 1.1<sup>*‡</sup> |
| AET (ms) | 67.6 ± 0.9 | 57.9 ± 1.3<sup>***‡</sup> | 77.2 ± 1.1<sup>#</sup> | 71.5 ± 2.0<sup>*‡</sup> |
| MPI | 0.68 ± 0.09 | 0.75 ± 0.07 | 0.54 ± 0.05<sup>\$</sup> | 0.76 ± 0.05<sup>*‡</sup> |
| E/E' | 24.2 ± 1.2 | 29.2 ± 1.5 | 33.3 ± 3.3 | 39.3 ± 3.5<sup>#</sup> |

LVID, left ventricular internal diameter; IVS, interventricular septal wall thickness; LVPW, LV posterior wall thickness; LVRWT, LV relative wall thickness; s, systole; d, diastole; EF, ejection fraction; FS, fractional shortening; IVCT, isovolumetric contraction time; IVRT, isovolumetric relaxation time; AET, aortic ejection time; MPI, myocardial performance index; E, early/passive filling velocity of LV; E', early mitral annulus velocity. All values are mean ± SEM as analyzed by 2-way ANOVA followed by Tukey's multiple comparison test. #- P<0.05 when compared to Sham WT, \$- P<0.05 when compared to Sham KO, ‡- P<0.05 when compared to WT control. In some cases, an unpaired Student's t-test was used to compare WT and KO groups for Sham or TAC treatments; \*- P<0.05, \*\*- P<0.005, \*\*\*- P<0.0005.

### 3.7 *Smtnl1*^-/-^ hearts display characteristics of diastolic dysfunction that are amplified following TAC

As detailed in **Table 3**, the hemodynamic parameters derived from the PV relationship determined after Sham surgery or 4 weeks of TAC revealed critical differences in measurements associated with diastolic function. The Sham KO mice display increased end diastolic pressure (EDP, 14 ± 1 mmHg [KO] vs. 6 ± 2 mmHg [WT], p = 0.007), end diastolic pressure volume relationship slope (EDPVR, 0.52 ± 0.05 [KO] vs 0.26 ± 0.03 [WT], p = 0.0004) and isovolumetric relaxation time constant (Tau, 10.98 ± 0.62 ms [KO] vs 8.34 ± 0.73 ms [WT], p =0.015). Interestingly, these diastolic parameters were not different between genotypes at 4 weeks post-TAC. In this case, significantly greater proportional change was observed for the EDP of the WT mice when compared to their Sham counterparts (**Figure 7D**; 10 ± 2 mmHg [WT] vs. 3 ± 2 mmHg [KO], p = 0.0249). Additionally, increased isovolumetric relaxation time (IVRT, 28.9 ± 1.1 ms [KO] vs. 24.1 ± 1.3 ms [WT], p = 0.044) and myocardial performance index (MPI, 0.758 ± 0.048 [KO] vs. 0.544 ± 0.050 [WT], p = 0.027) were observed when echocardiographic measurements of the TAC KO group were compared to the TAC WT group (**Table 2**). These observations suggest an amplified development of diastolic dysfunction under pressure overload in the *Smtnl1*^-/-^ mice. This is evident with substantial proportional increases in IVRT (**Figure 7E;** −2.45 ± 1.30 ms [WT] vs 2.31 ± 1.07 ms [KO], p = 0.019) and MPI (**Figure 7F**; −0.14 ± 0.06 [WT] vs 0.01 ± 0.05 [KO], p = 0.057) in TAC KO from their Sham controls as well as increased E/E’ ratio following TAC in the *Smtnl1*^-/-^ mice (39.3 ± 3.5 [TAC KO] vs 29.2 ± 1.5 [Sham KO], p = 0.018) (**Table 3**).

## 4. Discussion

Cardiac and vascular function are irrevocably intertwined, with changes in cardiac output driving changes in vascular contractility, and changes in vascular resistance driving changes in cardiac function (Levy and Pappano, 2007). Previous reports have revealed important impacts for SMTNL1 on cardiovascular performance; therefore, we characterized the structural and functional phenotype of the heart in the absence of SMTNL1, at baseline and under cardiac stress with pressure overload induced by constrictive banding of the aorta. The results of our study reveal that SMTNL1 plays a critical role in modulating cardiac functionality in addition to its previously reported impacts on vascular performance (Turner et al., 2014; Turner et al., 2019; Wooldridge et al., 2008; Lontay et al., 2010).

Male mice with global deficiency in SMTNL1 had smaller hearts but possessed a substantially larger LV mass in comparison to male WT mice, which was further reflected in the echocardiogram that revealed thicker LV posterior and anterior walls and smaller internal LV diameter during systole. Coincident with these morphometric alterations in heart structure, the circulating levels of pro-MMP9, a pro-hypertrophic biomarker (Halade et al., 2013), were elevated in KO mice. Sex dimorphism was also seen in the functional aspect with improvement in systolic performance as evidenced by increased EF% and FS% apparent only in the male *Smtnl1*^-/-^ mice. The corresponding observation of increased aortic and pulmonary arterial flow suggests that cardiac output was selectively enhanced in the *Smtnl1*^-/-^ male group. This novel finding is in agreement with past reports of sex dimorphism that linked SMTNL1 deletion with enhanced cardiovascular performance and an exercise-adapted vascular phenotype (Wooldridge et al., 2008). In this regard, male *Smtnl1*^-/-^ mice exhibited longer time to fatigue and greater distances run than male WT littermates when subjected to a forced-running exercise program. SMTNL1 deletion was also associated with enhanced myogenic reactivity of vessels isolated from male, but not female, mice (Turner et al., 2019). The augmented pressure-induced contractility of mesenteric resistance vessels from *Smtnl1*^-/-^ mice was linked to changes in protein kinase C signaling and the activity of myosin phosphatase. Herein, electrocardiographic findings in male *Smtnl1*^-/-^ animals revealed a reduction in resting heart rate with a notably lengthened RR interval, and an enhanced conductance as evident from the shortened P (atrial depolarization) and T (ventricular repolarization) wave duration and increased amplitude of S wave in the *Smtnl1*^-/-^ hearts. Slow resting heart rate is a physiological adaptation found with endurance-exercise that acts to maintain a normal cardiac output and blood pressure despite the training-induced increase in stroke volume (D’Souza et al., 2014). Thus, it is clear that SMTNL1 deletion dramatically affects cardiac structure and function as well as providing an aerobic exercise-adapted cardiovascular phenotype with reduced resting heart rate and enhanced electrical propagation (Gielen et al., 2010).

Aortic constriction was used to induce an abrupt pressure overload and steady mechanical stress to the heart (Tarnavski, 2009; Bosch et al., 2020). As expected, the systolic pressure of both genotypes was elevated under conditions of TAC-induced pressure overload. In addition, characteristic concentric LV remodeling along with marked interstitial fibrosis was observed for both genotypes. While both *Smtnl1*^-/-^ and WT developed similar end-points of cardiac hypertrophy, the sham KO mice initially had a smaller heart than its WT counterpart. Considering the above, the similar end-points in cardiac phenotype observed for TAC-challenged WT and KO hearts suggests the activation of adaptive mechanisms to maintain LV function (i.e., CO, EF% and FS%) under conditions of elevated basal LV hemodynamic load due to the vascular impact of SMTNL1 deletion. This was further corroborated by the echocardiographic M-mode measurements of näive KO mice that possessed significantly increased LV mass and relative wall thicknesses. Moreover, *Smtnl1*^-/-^ mice in the Sham group, had elevated systolic pressure and mean arterial pressure due to substantially augmented total peripheral resistance in the absence of any aortic constriction. The findings stress the importance of SMTNL1 deletion in advancing adverse cardiac responses to a hyperconstrictive vasculature. The splanchnic mesentery receives 10-40% of the cardiac output depending on digestion status (Harper and Chandler, 2016; Sun et al., 1992), and male *Smtnl1*^-/-^ mice possess significantly enhanced vasoconstriction of the mesenteric arteries that would contribute to elevated systemic vascular resistance (Turner et al., 2019). However, an earlier characterization of the *Smtnl1*^-/-^ animals is in disagreement since it did not reveal any alteration in peripheral blood pressure (Wooldridge et al., 2008). This discrepancy could be due to the differences in measurement techniques, or the age of the study animals. The Wooldridge report used non-invasive tail-cuff plethysmography on awake animals at an age of 8 weeks, while the current study employed a 1 F Millar pressure catheter inserted to the aorta with anesthetized animals at an age of 12-13 weeks

The systemic vascular resistance of the *Smtnl1*^-/-^ mice with TAC-burden further increased in comparison to the WT but not the *Smtnl1*^-/-^ mice in the Sham-operated group, showing that only the WT mice did trend to develop a higher vascular resistance upon pressure overload. This implies that the *Smtnl1*^-/-^ animals were more adapted to withstand the acute pressure overload inflicted by the aortic banding than their WT counterparts, presumably due to the inherent presence of enhanced myogenic reactivity of the microvasculature (Turner & MacDonald, 2014; Turner et al., 2019). Interestingly, PKG-Iα is reported to inhibit cardiac remodeling in TAC-challenged hearts (Blanton et al 2012). The PKG-dependent phosphorylation of SMTNL1 is also known to modulate its cellular function (Borman et al., 2004; Wooldridge et al., 2008; Borman et al., 2009; Lontay et al., 2010), so whether SMTNL1 is integrated into this PKG-Iα protective response warrants further investigation. Like SMTNL1, silencing of other vasoregulatory substrates of PKG-Iα has been reported to cause hypertension, including IRAG (Geiselhöringer et al., 2004), RGS2 (Tang et al., 2003), and the BKCa channel (Pluger et al., 2000).

The morphological changes of the hearts after acute pressure overload were accompanied by a preserved systolic function in both genotypes. Indeed, the load-dependent systolic parameters ejection fraction, shortening, cardiac output, stroke work, stroke volume and maximum (+) dP/dt as well as the load-independent parameters viz., preload recruited stroke work and end systolic pressure volume relationship (i.e., systolic elastance) were not different between genotypes in both sham and TAC groups. These findings support normal systolic pump performance of the LV under conditions of SMTNL1 KO with or without the acute pressure overload. The introduction of cardiac stress in the form of acute pressure overload also caused concentric hypertrophy and fibrotic deposition to develop without LV dilation in both genotypes. Although not observed in our study, ventricular cavity expansion due to a decline in EF% and developing systolic heart failure is a typical cardiac outcome of TAC banding (Mohammed et al., 2012; Weinheimer et al., 2015). Progression to heart failure does depend on the severity and duration of the aortic constriction, and other reports have suggested that significant pathologic LV dilatation is not observed during the first 4 weeks of pressure-overload following TAC, due to modest systolic dysfunction and fibrotic accumulation (Zankov et al., 2017).

A key piece of novel evidence that has emerged from our studies is the presence of impaired diastolic function in the *Smtnl1*^-/-^ cohort. Hemodynamic measurements employing a pressure-conductance catheter system, the most accurate technique for characterizing diastolic function (Burkhoff et al., 2005), revealed relaxation impairment in the Sham group of *Smtnl1*^-/-^ animals. These animals exhibited an increased time constant of isovolumetric relaxation (i.e., tau, τ), suggesting alterations in the timing of LV pressure fall during myocardial relaxation. Furthermore, elevated diastolic chamber stiffness was observed as evidenced by substantial increases in the end diastolic pressure and the preload-independent end diastolic pressure volume relationship as well as a trend toward elevated LV filling pressure due to an increase in E/E’ ratio. Diastolic dysfunction was exacerbated in *Smtnl1*^-/-^ mice with aortic banding as indicated by an increased E/E’ ratio when compared to the Sham group as well as a prolonged isovolumetric relaxation time and enhanced myocardial performance index when compared to the WT counterparts. In contrast, hemodynamic parameters such as tau, end diastolic pressure and end diastolic pressure volume relationship did not show a difference between the WT and *Smtnl1*^-/-^ cohorts after aortic banding. In this case, the WT mice tended to mimic the diastolic dysfunction observed in the Sham KO mice. The small exacerbation of diastolic dysfunction upon pressure overload in the KO animal relative to the response of WT animals suggests that it is the vascular effect of SMTNL1 that is responsible for the cardiac phenotype rather than altered cardiac function driving subsequent vascular adaptations.

*Smtnl1* expression was low in heart tissue, so it is unlikely that the cardiomyocyte is the primary effector cell targeted by SMTNL1 silencing. Additionally, SMTNL1 expression is not limited to the vascular smooth muscle (Wooldridge et al., 2008), and the use of a global *Smtnl1*^-/-^ animal has the potential to combine the local effect of SMTNL1 absence in the vasculature with systematic effects of SMTNL1 deletion in other tissue beds to provide extrinsic impact on the heart. SMTNL1 has its highest expression in the skeletal muscle (Wooldridge *et al*., 2008). This tissue, which makes up 30 to 40% of the body mass in healthy individuals, has recently become the focus of studies investigating its role as a secretory organ (Walsh, 2009; Norheim et al., 2011; Hamrick, 2012; Aoi, 2017; Giudice and Taylor, 2017). Muscle-derived secretory proteins or “myokines” can exert protective effects on the cardiovascular system. Interestingly, follistatin-like 1 (Fstl1) expression is upregulated in response to cardiac stress such as TAC-induced pressure overload (Oshima et al., 2008), and this myokine can act on vascular endothelial cells in an eNOS-dependent manner to promote function and survival (Ouchi et al, 2008). In addition, transcriptome profiling found apelin to be upregulated in human skeletal muscle with endurance-exercise training (Besse-Patin et al., 2014). Apelin is another myokine that is implicated in cardiovascular homeostasis with defined effects on vasodilation and myocardial contractility through its action on the endothelium (Zhong et al., 2017).

Stiffening of the vascular wall may occur from a loss of smooth muscle tone (Joannides et al., 2001) as well as from a reduction in nitric oxide production from endothelial cells (Bellien et al., 2010). Both Sham-operated and pressure overloaded *Smtnl1*^-/-^ mice displayed elevated arterial elastance (Ea) in comparison to the sham WT group. Load-dependent systolic parameters and LV pump performance, including EF, FS, cardiac output, stroke work, stroke volume and maximum (+) dP/dt as well as the load-independent parameters viz., preload recruited stroke work and end systolic pressure volume relationship (systolic elastance) were not different between genotypes in both Sham and TAC groups. So, the findings again suggest that the relaxation impairment of *Smtnl1*^-/-^ hearts could be mainly due to changes in the vasculature rather than in the left ventricle. Ea is also increased in patients with diastolic heart failure, and it has been suggested that such vascular stiffening can contribute to the pathogenesis of diastolic dysfunction (Kawaguchi et al., 2003; Gaasch and Little, 2007). Furthermore, diastolic dysfunction is predominant in patients displaying heart failure with preserved ejection fraction (HFpEF) (Paulus & Tschope, 2013). Currently, the most widely proposed HFpEF paradigm links vascular impairment to diastolic dysfunction and LV remodeling (Kishimoto et al., 2017; Shantsila et al., 2012; Marechaux et al., 2016; Lee et al., 2016; Giamouzis et al., 2016). The suggested site of impact is the microvasculature, where systemic inflammatory effects driven by risk factors and disease modifiers induce endothelial dysfunction. This, in turn, elicits myocardial remodeling to result in compromised diastolic ventricular function that typically defines the HFpEF syndrome. Given that SMTNL1 silencing drives vascular stiffness and changes in serum biomarker profiles reflective of an injured or inflamed endothelium (i.e., sE-selectin, sP-selectin, pro-MMP9 and VEGF; Mozos et al., 2017) with subsequent development of diastolic dysfunction, further research to examine the impact of SMTNL1 on endothelial performance and the progressive evolution of HFpEF-like pathophysiology is warranted. The molecular pathways (and their sex dependency) acting on the endothelium to promote HFpEF development are poorly understood, and this limits development of effective therapeutic strategies that target endothelial dysfunction in order to ameliorate cardiovascular disease outcomes.

In conclusion, these data highlight the emergence of SMTNL1 as a novel contributor to cardiac performance. We now demonstrate that global silencing of SMTNL1 in mice is associated with changes in cardiac structure to promote the nascent development of diastolic dysfunction with impaired LV relaxation and enhanced passive stiffness in a sex-dimorphic way. The emergence of diastolic dysfunction in young *Smtnl1*^-/-^ animals is likely due to elevations in systemic vascular resistance and arterial stiffness. Future studies should investigate the influence of SMTNL1 on the endothelium as well as its impact on the release of cardioprotective myokines from skeletal muscle. Collectively, these findings reveal novel importance for SMTNL1 in the pathogenesis of vascular dysfunction, compromised diastolic ventricular dysfunction, and the development of a HFpEF-like syndrome in mice.

## Acknowledgements

This work was supported by a research grant from the Canadian Institutes of Health Research (MOP#97931 to J.A.M.). S.R.T. was recipient by an Alberta Innovates Health Solutions (AIHS) Doctoral Studentship and a CIHR Fredrick Banting and Charles Best Canada Graduate Scholarship. M.M. was supported by an Achievers in Medicine Doctoral Scholarship from the University of Calgary. W.C.C. is holder of the Andrew Family Professorship at the University of Calgary. The authors gratefully acknowledge Dr. Yong-Xiang Chen and the Libin Institute Histopathology Core for assistance with cardiac tissue sectioning and staining.

## Author Contributions

M.M. and S.R.T. designed, performed and analyzed the experiments. D.B. assisted with echocardiography, PV loops, and TAC surgeries. M.M. and J.A.M. wrote the manuscript; M.M., S.R.T., and J.A.M. prepared figures. W.C.C. and J.A.M. edited the manuscript, supervised trainees and provided intellectual contributions to the project. J.A.M. conceived and coordinated the study. All authors reviewed the results and approved the final version of the manuscript.

## Authors’ Competing Interests Statement

J.A.M. is cofounder and has an equity position in Arch Biopartners Inc. All other authors declare no conflicts of interest.

## Notes

### Competing Interest Statement

Justin A. MacDonald is cofounder and has an equity position in Arch Biopartners Inc. All other authors declare no conflicts of interest.

